# Combining global tree cover loss data with historical national forest-cover maps to look at six decades of deforestation and forest fragmentation in Madagascar

**DOI:** 10.1101/147827

**Authors:** Ghislain Vieilledent, Clovis Grinand, Fety A. Rakotomalala, Rija Ranaivosoa, Jean-Roger Rakotoarijaona, Thomas F. Allnutt, Frédéric Achard

**Author notes:** Corresponding author: \ \.

## Abstract

The island of Madagascar has a unique biodiversity, mainly located in the tropical forests of the island. This biodiversity is highly threatened by anthropogenic deforestation. Existing historical forest maps at national level are scattered and have substantial gaps which prevent an exhaustive assessment of long-term deforestation trends in Madagascar. In this study, we combined historical national forest cover maps (covering the period 1953-2000) with a recent global annual tree cover loss dataset (2001-2014) to look at six decades of deforestation and forest fragmentation in Madagascar (from 1953 to 2014). We produced new forest cover maps at 30 m resolution for the year 1990 and annually from 2000 to 2014 over the full territory of Madagascar. We estimated that Madagascar has lost 44% of its natural forest cover over the period 1953-2014 (including 37% over the period 1973-2014). Natural forests cover 8.9 Mha in 2014 (15% of the national territory) and include 4.4 Mha (50%) of moist forests, 2.6 Mha (29%) of dry forests, 1.7 Mha of spiny forests (19%) and 177,000 ha (2%) of mangroves. Since 2005, the annual deforestation rate has progressively increased in Madagascar to reach 99,000 ha/yr during 2010-2014 (corresponding to a rate of 1.1%/yr). Around half of the forest (46%) is now located at less than 100 m from the forest edge. Our approach could be replicated to other developing countries with tropical forest. Accurate forest cover change maps can be used to assess the effectiveness of past and current conservation programs and implement new strategies for the future. In particular, forest maps and estimates can be used in the REDD+ framework which aims at “Reducing Emissions from Deforestation and forest Degradation” and for optimizing the current protected area network.

## 1 Introduction

Separated from the African continent and the Indian plate about 165 and 88 million years ago respectively (Ali & Aitchison, 2008), the flora and fauna of Madagascar followed its own evolutionary path. Isolation combined with a high number of micro-habitats (Pearson & Raxworthy, 2009) has led to Madagascar’s exceptional biodiversity both in term of number of species and endemism in many taxonomic groups (Crottini *et al*., 2012; Goodman & Benstead, 2005). Most of the biodiversity in Madagascar is concentrated in the tropical forests of the island which can be divided into four types: the moist forest in the East, the dry forest in the West, the spiny forest in the South and the mangroves on the West coast (Vieilledent *et al*., 2016). This unparalleled biodiversity is severely threatened by deforestation (Harper *et al*., 2007; Vieilledent *et al*., 2013) associated with human activities such as slash-and-burn agriculture and pasture (Scales, 2011). Tropical forests in Madagascar also store a large amount of carbon (136 MgC.ha^−1^ in the moist forest, Vieilledent *et al*., 2016) and high rates of deforestation in Madagascar (1.4–4.7 %/yr, Achard *et al*., 2002) are responsible for large CO_2_ emissions in the atmosphere. Deforestation threatens species survival by directly reducing their available habitat (Brooks *et al*., 2002; Tidd *et al*., 2001). Forest fragmentation can also lead to species extinction by isolating populations from each other and creating forest patches too small to maintain viable populations (Saunders *et al*., 1991). Fragmentation also increases forest edge where ecological conditions (such as air temperature, light intensity and air moisture) can be dramatically modified, with consequences on the abundance and distribution of species (Broadbent *et al*., 2008; Gibson *et al*., 2013; Murcia, 1995). Forest fragmentation can also have substantial effects on forest carbon storage capacity, as carbon stocks are about 50% lower at the forest edge than under a closed canopy (Brinck *et al*., 2017). Moreover, forest carbon stocks vary spatially due to climate or soil factors (Saatchi *et al*., 2011; Vieilledent *et al*., 2016). As a consequence, accurate and spatially explicit maps of forest cover and forest cover change are necessary to monitor biodiversity loss and carbon emissions from deforestation and forest fragmentation, assess the efficiency of present conservation strategies (Eklund *et al*., 2016), and implement new strategies for the future (Vieilledent *et al*., 2016, 2013). Simple time-series of forest cover estimates, such as those provided by the FAO Forest Resource Assessment report (Keenan *et al*., 2015) are not sufficient.

Unfortunately, accurate and exhaustive forest cover maps are not available for Madagascar after year 2000. Harper *et al*. (2007) produced maps of forest cover and forest cover changes over Madagascar for the years 1953, 1973, 1990 and 2000. The 1953 forest map is a vector map derived from the visual interpretation of aerial photographs. Forest maps for the years 1973, 1990, and 2000 were obtained from the supervised classification of Landsat satellite images and can be used to derive more accurate estimates of forest cover than those from the FAO Forest Resource Assessment report. Nonetheless, maps provided by Harper *et al*. (2007) are not exhaustive (due to the presence of clouds in the satellite imagery), e.g. 11 244 km2 are mapped as unknown cover type for the year 2000. Using a similar supervised classification approach as in Harper *et al*. (2007), more recent maps have been produced for the periods 2000-2005-2010 by national institutions, with the technical support of international environmental NGOs (MEFT *et al*., 2009; ONE *et al*., 2013). Another set of recent forest cover maps using an advanced statistical tool for classification, the Random Forest classifier (Grinand *et al*., 2013; Rakotomala *et al*., 2015), was produced for the periods 2005-2010-2013 (ONE *et al*., 2015). However, these maps are either too old to give recent estimates of deforestation (MEFT *et al*., 2009; ONE *et al*., 2013), include large areas of missing information due to images with high percentage of cloud cover (ONE *et al*., 2013), or show large mis-classification in specific areas, especially in the dry and spiny forest domain, for which the spectral signal shows strong seasonal variations due to the deciduousness of such forests (overall accuracy is lower than 0.8 for the dry and spiny forests for the maps produced by ONE *et al*. (2015)). Moreover, the production of such forest maps from a supervised classification approach requires significant resources, especially regarding the image selection step (required to minimize cloud cover) and the training step (visual interpretation of a large number of polygons needed to train the classification algorithm) (Rakotomala *et al*., 2015). Most of this work of image selection and visual interpretation would need to be repeated to produce new forest maps in the future using a similar approach.

Global forest or tree cover products have also been published recently and can be tested at the national scale for Madagascar. Kim *et al*. (2014) produced a global forest cover change map from 1990 to 2000 (derived from Landsat imagery). This product was updated to cover the period 1975-2005 (http://glcf.umd.edu/data/landsatFCC/) but forest cover maps after 2005 were not produced. Moreover, the approach used in *Kim et al*. (2014) did not accurately map the forests in the dry and spiny ecosystems of Madagascar (see Fig. 8 in Kim *et al*. 2014). Hansen *et al*. (2013) mapped tree cover percentage, annual tree cover loss and gain from 2000 to 2012 at global scale at 30 m resolution. This product has since been updated and is now available up to the year 2014 (Hansen *et al*., 2013). To map forest cover from the Hansen *et al*. (2013) product, a tree cover threshold must be selected (that defines forest cover). Selecting such a threshold is not straightforward as the accuracy of the global tree cover map strongly varies between forest types, and is substantially lower for dry forests than for moist forests (Bastin *et al*., 2017). Moreover, the Hansen *et al*. (2013) product does not provide information on land-use. In particular the global tree cover map does not separate tree plantations such as oil palm or eucalyptus plantations from natural forests (Tropek *et al*., 2014). Thus, the global tree cover map from Hansen *et al*. (2013) cannot be used alone to produce a map of forest cover (Tyukavina *et al*., 2017).

In this study, we present a simple approach which combines the historical forest maps from Harper *et al*. (2007) and more recent global products from Hansen *et al*. (2013) to derive annual wall-to-wall forest cover change maps over the period 2000-2014 for Madagascar. We use the forest cover map provided by Harper *et al*. (2007) for the year 2000 (defining the land-use) with the tree cover loss product provided by Hansen *et al*. (2013) that we apply only inside forest areas identified by Harper *et al*. (2007). Similar to the approach of Harper *et al*. (2007), we also assess trends in deforestation rates and forest fragmentation from 1953 to 2014. We finally discuss the possibility to extend our approach to other tropical countries or repeat it in the future for Madagascar. We also discuss how our results could help assess the effectiveness of past and current conservation strategies in Madagascar, and implement new strategies in the future.

## 2 Materials and Methods

### 2.1 Creation of new forest cover maps of Madagascar from 1953 to 2014

Original 1990-2000 forest cover change map for Madagascar from Harper *et al*. (2007) is a raster map at 28.5 m resolution. It was derived from the supervised classification of Landsat TM (Thematic Mapper) and ETM+ (Enhanced Thematic Mapper Plus) satellite images. For our study, this map has been resampled at 30 m resolution using a nearest-neighbor interpolation and reprojected in the WGS 84/UTM zone 38S projected coordinate system.

The 2000 Harper’s forest map includes 208,000 ha of unclassified areas due to the presence of clouds on satellite images. Unclassified areas were mostly (88%) present within the moist forest domain which covered 4.17 Mha in 2000. To provide a label (forest or non-forest) to these unclassified pixels, we used the 2000 tree cover percentage map of Hansen *et al*. (2013) and selected a tree cover threshold of 75% to define the forest (Achard *et al*., 2014; Aleman *et al*., 2017). This threshold allows to characterize properly the moist forest in Madagascar as 90% of the moist forest in 2000 in Harper *et al*. (2007) has a tree cover greater than 75% (Fig. A1). For this step, the Hansen’s 2000 tree cover map was resampled on the same grid as the original Harper’s map at 30 m resolution using a bilinear interpolation. We thus obtained a forest cover map for the year 2000 covering the full territory of Madagascar.

We then combined the forest cover map of the year 2000 with the annual tree cover loss maps from 2001 to 2014 from Hansen *et al*. (2013) to create annual forest cover maps from 2001 to 2014 at 30 m resolution. To do so, Hansen’s tree cover loss maps were resampled on the same grid as the original Harper’s map at 30 m resolution using a nearest-neighbor interpolation. We also completed the Harper’s forest map of year 1990 by filling unclassified areas (due to the presence of clouds on satellite images) using our forest cover map of year 2000. To do so, we assumed that if forest was present in 2000, the pixel was also forested in 1990. Indeed, there is little evidence of natural forest regeneration in Madagascar (Grouzis *et al*., 2001; Harper *et al*., 2007), especially over such a short period of time. The remaining unclassified pixels were limited to a relatively small total area of about 8,000 ha. We labeled these residual pixels as non-forest, as for the year 2000.

The 1973 forest cover map for Madagascar from Harper *et al*. (2007) is a raster map at 57 m resolution derived from the supervised classification of Landsat MSS (Multispectral Scanner System) satellite images. We resampled this map at 30 m resolution using a nearest-neighbor interpolation on the same grid as the forest cover maps for years 1990 and 2000. We completed the Harper’s forest map of year 1973 by filling unclassified areas using our forest cover map of the year 1990 assuming that if forest was present in 1990, it was also present in 1973. Contrary to the year 1990, the remaining unclassified pixels for year 1973 corresponded to a significant total area of 3.3 millions ha which was left as is.

The 1953 forest cover map from Harper *et al*. (2007) is a vector map produced by scanning the paper map of Humbert *et al*. (1965) which was derived from the visual interpretation of aerial photographs. We reprojected the forest cover map of year 1953 in the WGS 84/UTM zone 38S projected coordinate system. Because of the methodology used to derive the 1953 forest cover map, it was not possible to perfectly aligned this map with the forest cover maps of later years which were produced through digital processing of satellite imagery. As a consequence, the 1953 cannot be merged with the map of later years to identify precisely the location of the deforested areas. Nonetheless, the 1953 forest cover map can be used to have a rough estimate of the forest cover and forest fragmentation at this date. To do so, the map was rasterized at 30 m resolution on the same grid as the forest cover maps for years 1973, 1990 and 2000.

Finally for all forest cover maps from 1973, isolated single non-forest pixels (i.e. fully surrounded by forest pixels) were recategorized as forest pixels. Doing so, forest cover increased from about 95,000 ha for year 1953 to about 600,000 ha for year 2010. This allowed us to avoid counting very small scale events (*<*0.1 ha, such as selective logging or wind-throw) as deforestation. It also prevents us from underestimating forest cover and overestimating forest fragmentation.

### 2.2 Computing forest cover areas and deforestation rates

From these new forest cover maps, we calculated the total forest cover area for seven available years (1953-1973-1990-2000-2005-2010-2014), and the annual deforested area and annual deforestation rate for the corresponding six time periods between 1953 and 2014. The annual deforestation rates were calculated using Eq. 1 (Puyravaud, 2003; Vieilledent *et al*., 2013):

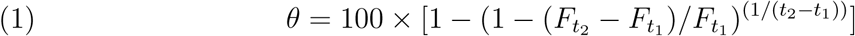

In Eq. 1, *θ* is the annual deforestation rate (in %/yr), *F*_*t*_2__ and *F*_*t*_1__ are the forest cover free of clouds at both dates *t*_2_ and *t*_1_, and *t*_2_ *− t*_1_ is the time-interval (in years) between the two dates.

Because of the large unclassified area (3.3 millions ha) in 1973, the annual deforestation areas and rates for the two periods 1953-1973 and 1973-1990 are only partial estimates computed on the basis of the available forest extent. Area and rate estimates are produced at the national scale and for the four forest types present in Madagascar: moist forest in the East, dry forest in the West, spiny forest in the South, and mangroves on the Western coast (Fig. 1). To define the forest types, we used a map from the MEFT (*“Ministère de l’Environnement et des Forêts à Madagascar”*) with the boundaries of the four ecoregions in Madagascar. Ecoregions were defined on the basis of climatic and vegetation criteria using the climate classification by Cornet (1974) and the vegetation classification from the 1996 IEFN national forest inventory (Ministère de l’Environnement, 1996). Because mangrove forests are highly dynamic ecosystems that can expand or contract on decadal scales depending on changes in environmental factors (Armitage *et al*., 2015), a fixed delimitation of the mangrove ecoregion on six decades might not be fully appropriate. As a consequence, our estimates of the forest cover and deforestation rates for mangroves in Madagascar must be considered with this limitation.

**Figure 1:**
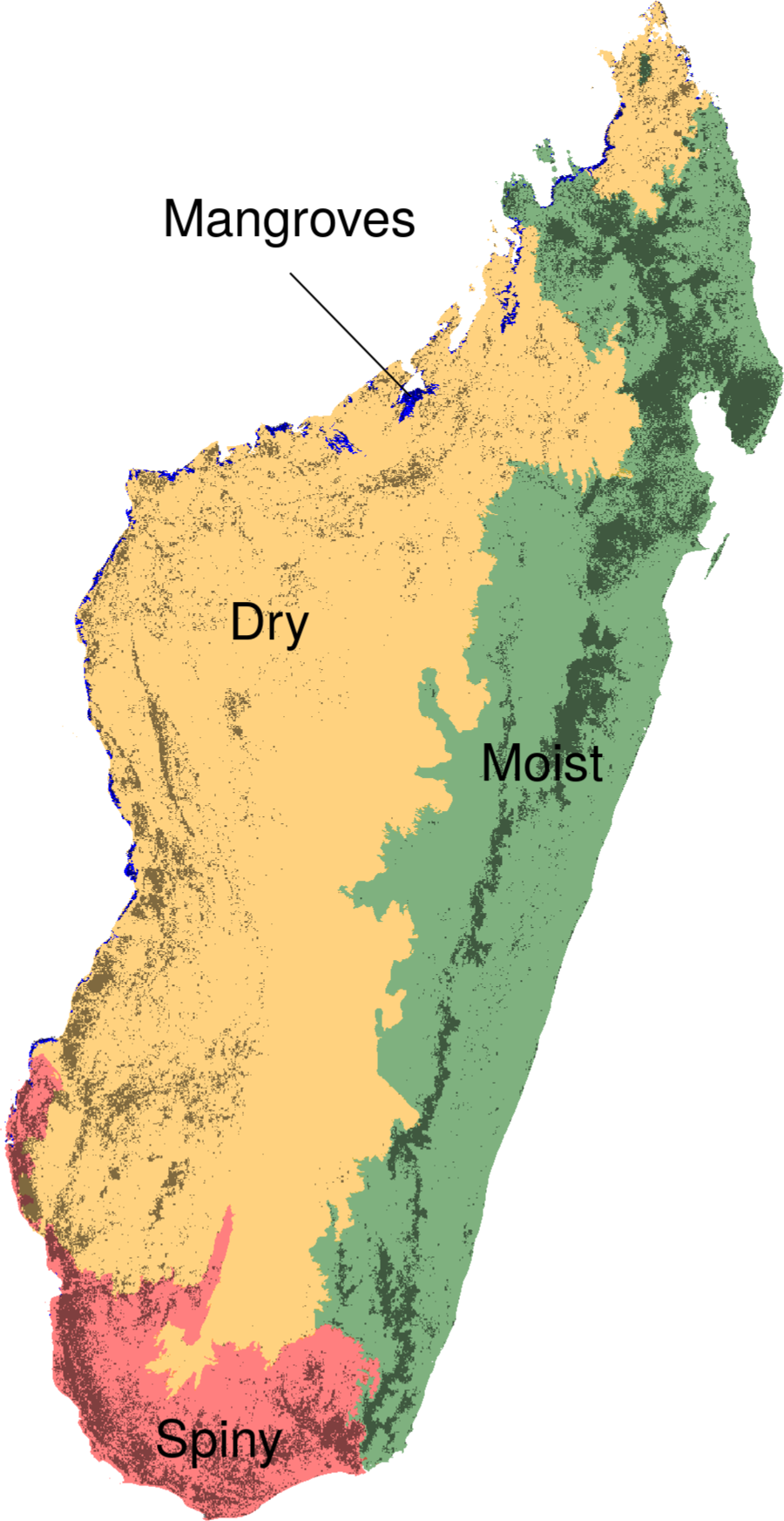
Ecoregions and forest types in Madagascar. Madagascar can be divided into four climatic ecoregions with four forest types: the moist forest in the East (green), the dry forest in the West (orange), the spiny forest in the South (red), and the mangroves on the West coast (blue). Ecoregions were defined following climatic (Cornet, 1974) and vegetation (Ministère de l’Environnement, 1996) criteria. The dark grey areas represent the remaining natural forest cover for the year 2014. Forest types are defined on the basis of their belonging to one of the four ecoregions.

### 2.3 Comparing our forest cover and deforestation rate estimates with previous studies

We compared our estimates of forest cover and deforestation rates with estimates from the three existing studies at the national scale for Madagascar: (i) Harper *et al*. (2007), (ii) MEFT *et al*. (2009) and (iii) ONE *et al*. (2015). Harper *et al*. (2007) provides forest cover and deforestation estimates for the periods c. 1953-c. 1973-1990-2000. *MEFT et al*. (2009) provides estimates for the periods 1990-2000-2005 and ONE *et al*. (2015) provides estimates for the periods 2005-2010-2013. To compare our forest cover and deforestation estimates over the same time periods, we consider an additional time-period in our study (2010-2013) by creating an extra forest cover map for the year 2013. We computed the Pearson’s correlation coefficient and the root mean square error (RMSE) between our forest cover estimates and forest cover estimates from previous studies for all the dates and forest types (including also the total forest cover estimates). For previous studies, the computation of annual deforestation rates (in %/yr) is not always detailed and might slightly differ from one study to another (see Puyravaud, 2003). Harper *et al*. (2007) also provide total deforested areas for the two periods 1973-1990 and 1990-2000. We converted these values into annual deforested area estimates. When annual deforested areas were not reported (for 1953-1973 in Harper *et al*. (2007) and in MEFT *et al*. (2009) and *ONE et al*. (2015)), we computed them from the forest cover estimates in each study. These estimates cannot be corrected from the potential bias due to the presence of residual clouds. Forest cover and deforestation rates were then compared between all studies for the whole of Madagascar and the four ecoregions. The same ecoregion boundaries as in our study were used in ONE *et al*. (2015) but this was not the case for Harper *et al*. (2007) and MEFT *et al*. (2009), which can explain a part of the differences between the estimates.

### 2.4 Fragmentation

We also conducted an analysis of changes in forest fragmentation for the years 1953, 1973, 1990, 2000, 2005, 2010 and 2014 at 30 m resolution. We used a moving window of 51 *×* 51 pixels (corresponding to an area of about 2.34 km^2^) centered on each forest pixel to compute the percentage of forest pixels in the neighborhood. We used this percentage as an indication of the forest fragmentation (Riitters & Wickham, 2012; Vogt & Riitters, 2017). The size of the moving windows was based on a compromise: a sufficiently high number of cells (here 2601) had to be considered to be able to compute a percentage and a reasonably low number of cells had to be chosen to have a local estimate of the fragmentation. Water bodies were not masked when computing the percentage of forest pixels, meaning that forest located near a water body was considered as fragmented. Computations were done using the function r.neighbors of the GRASS GIS software (Neteler & Mitasova, 2008). Using the density of forest in the neighborhood, we defined five forest fragmentation classes: 0-20% (highly fragmented), 21-40%, 41-60%, 61-80% and 81-100% (lowly fragmented). We reported the percentage of forest falling in each fragmentation class for the six years and analyzed the dynamics of fragmentation over the six decades.

We also computed the distance to forest edge for all forest pixels for the years 1953, 1973, 1990, 2000, 2005, 2010 and 2014. For that, we used the function `gdal proximity.py` of the GDAL library (http://www.gdal.org/). We computed the mean and 90% quantiles (5% and 95%) of the distance to forest edge and looked at the variation of these values over time. Previous studies have shown that forest micro-habitats were mainly altered within the first 100 m of the forest edge (Brinck *et al*., 2017; Broadbent *et al*., 2008; Murcia, 1995). Consequently, we also estimated the percentage of forest within the first 100 m of the forest edge for each year and looked at the variation of this percentage over the six decades.

## 3 Results

### 3.1 Forest cover change and deforestation rates

Natural forests in Madagascar covered 16.0 Mha in 1953, about 27% of the national territory of 587,041 km2. In 2014, the forest cover dropped to 8.9 Mha, corresponding to about 15% of the national territory (Fig. 2 and Tab. 1). Madagascar has lost 44% of its natural forest between 1953 and 2014, including 37% between 1973 and 2014 (Fig. 2 and Tab. 1). In 2014 the remaining 8.9 Mha of natural forest were distributed as follow: 4.4 Mha of moist forest (50% of total forest cover), 2.6 Mha of dry forest (29%), 1.7 Mha of spiny forest (19%) and 0.18 Mha (2%) of mangrove forest (Fig. 1 and Tab. 2).

**Figure 2:**
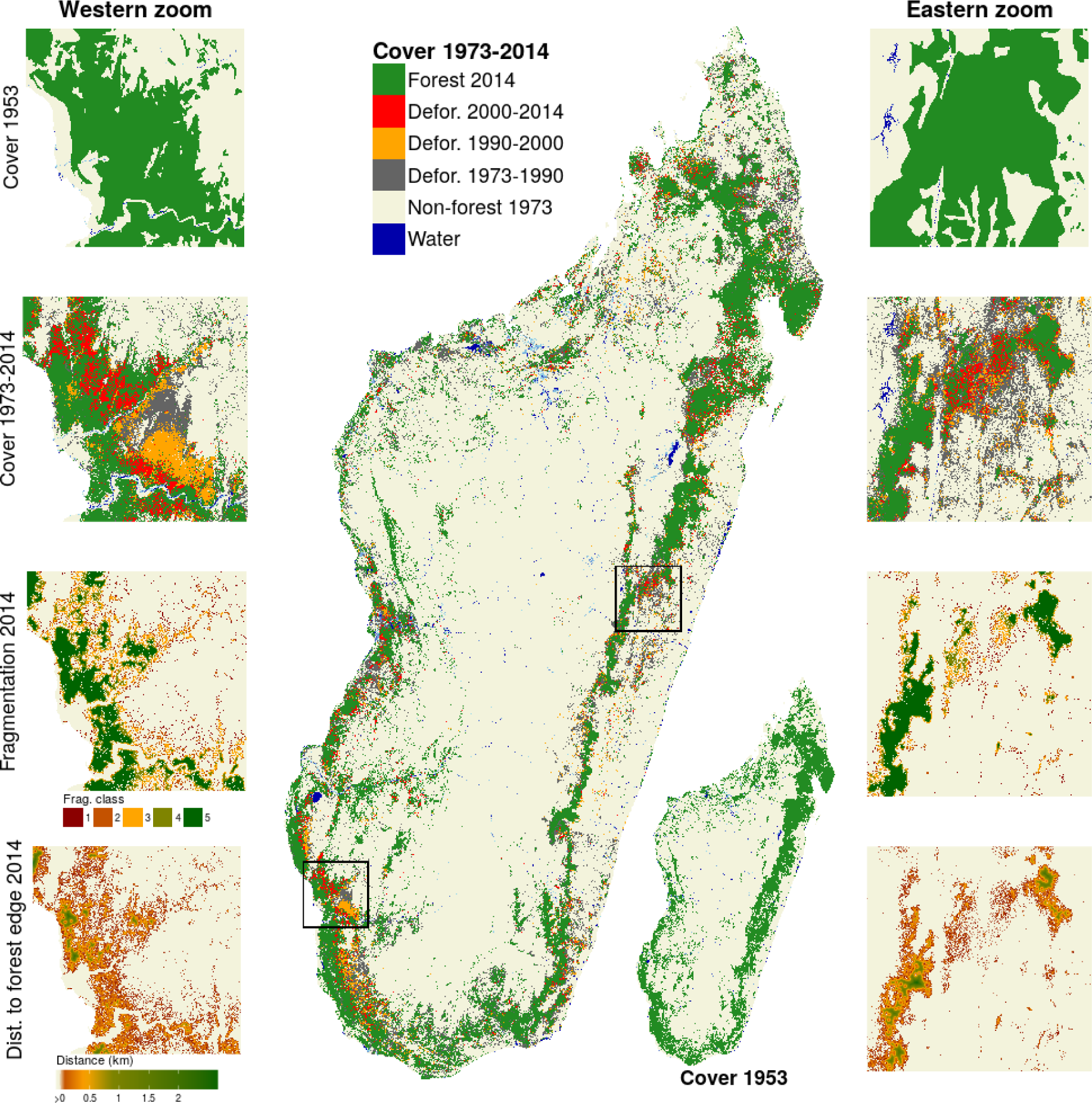
Forest cover change on six decades from 1953 to 2014 in Madagascar. forest cover changes from 1973 to 2014 are shown in the main figure, and forest cover in 1953 is shown in the bottom-right inset. Two zooms in the western dry (left part) and eastern moist (right part) ecoregions present more detailed views of (from top to bottom): forest cover in 1953, forest cover change from 1973 to 2014, forest fragmentation in 2014 and distance to forest edge in 2014. Data on water bodies (blue) and water seasonality (light blue for seasonal water to dark blue for permanent water) has been extracted from Pekel *et al*. (2016).

**Table 1:** Change in natural forest cover and deforestation rates from 1953 to 2014 in Madagascar. Areas are provided in thousands of hectares (Kha). Forest map for the year 1973 has 3.3 Mha of unclassified areas due to the presence of clouds on satellite images. As a consequence, partial deforestation rates for the periods 1953-1973 and 1973-1990 are computed based on the available forest extent. The last two columns indicate the annual deforested areas and annual deforestation rates on the previous time-period (e.g. 1953-1973 for year 1973, 1973-1990 for year 1990, etc.).

| Year | Forest (Kha) | Unmap (Kha) | Annual defor. (Kha/yr) | Rate (%/yr) |
| --- | --- | --- | --- | --- |
| 1953 | 15,968 | 0 | - | - |
| 1973 | 14,243 | 3,317 | 86 | 0.6 |
| 1990 | 10,762 | 0 | 205 | 1.6 |
| 2000 | 9,879 | 0 | 88 | 0.8 |
| 2005 | 9,668 | 0 | 42 | 0.4 |
| 2010 | 9,320 | 0 | 70 | 0.7 |
| 2014 | 8,925 | 0 | 99 | 1.1 |

**Table 2:** Comparing Madagascar forest cover estimates with previous studies on the period 1953-2014. We compared our estimates of forest cover with the estimates from three previous studies (Harper *et al*., 2007; MEFT *et al*., 2009; ONE *et al*., 2015). Areas are provided in thousands of hectares (Kha). We obtained a Pearson’s correlation coefficient of 0.99 between our forest cover estimates and forest cover estimates from previous studies. The increase in mangrove and spiny forest covers from 1953 to 1973 in Harper *et al*. (2007) and our study is most probably due to differences in forest definition and mapping methods between the 1953 aerial-photography derived map and the 1973 Landsat image derived map.

| Forest type | Source | 1953 | 1973 | 1990 | 2000 | 2005 | 2010 | 2013 | 2014 |
| --- | --- | --- | --- | --- | --- | --- | --- | --- | --- |
| Total | Harper2007 | 15,996 | 14,173 | 10,606 | 8,982 | - | - | - | - |
|  | MEFT2009 | - | - | 10,650 | 9,678 | 9,413 | - | - | - |
|  | ONE2015 | - | - | - | - | 9,451 | 8,977 | 8,486 | - |
|  | this study | 15,968 | 14,243 | 10,762 | 9,879 | 9,668 | 9,320 | 9,051 | 8,925 |
| Moist | Harper2007 | 8,766 | 6,876 | 5,234 | 4,167 | - | - | - | - |
|  | MEFT2009 | - | - | 5,271 | 4,788 | 4,700 | - | - | - |
|  | ONE2015 | - | - | - | - | 4,556 | 4,457 | 4,345 | - |
|  | this study | 8,578 | 6,990 | 5,270 | 4,872 | 4,768 | 4,633 | 4,470 | 4,410 |
| Dry | Harper2007 | 4,252 | 4,028 | 2,712 | 2,457 | - | - | - | - |
|  | MEFT2009 | - | - | 3,321 | 3,085 | 3,028 | - | - | - |
|  | ONE2015 | - | - | - | - | 3,223 | 2,970 | 2,679 | - |
|  | this study | 4,762 | 4,435 | 3,225 | 2,941 | 2,881 | 2,735 | 2,642 | 2,596 |
| Spiny | Harper2007 | 2,978 | 3,030 | 2,420 | 2,132 | - | - | - | - |
|  | MEFT2009 | - | - | 2,124 | 1,872 | 1,757 | - | - | - |
|  | ONE2015 | - | - | - | - | 1,682 | 1,559 | 1,467 | - |
|  | this study | 2,463 | 2,583 | 2,055 | 1,858 | 1,811 | 1,744 | 1,731 | 1,713 |
| Mangroves | Harper2007 | - | - | 240 | 226 | - | - | - | - |
|  | MEFT2009 | - | - | - | - | - | - | - | - |
|  | ONE2015 | - | - | - | - | 174 | 171 | 170 | - |
|  | this study | 143 | 200 | 181 | 178 | 177 | 177 | 177 | 177 |

The forest cover change map produced on the period 1953-2014 (Fig. 2) allows to identify hot-spots of deforestation. Among the many recent hot-spots of deforestation visible on the map for the period 2000-2014, one is located at the south of the CAZ (*“Corridor Ankeniheny Zahamena”*) protected area, in the moist forest at the east of Madagascar (see eastern zoom in Fig. 2). Another major hot-spot of deforestation is located around the Ranobe-PK32 new protected area, in the dry forest at the south-west of Madagascar (see western zoom in Fig. 2).

Regarding the deforestation trend, we observed a progressive decrease of the deforestation rate after 1990 from 205,000 ha/yr (1.6%/yr) over the period 1973-1990 to 42,000 ha/yr (0.4%/yr) over the period 2000-2005 (Tab. 1). Then from 2005, the deforestation rate has progressively increased and has more than doubled over the period 2010-2014 (99,000 ha/yr, 1.1%/yr) compared to 2000-2005 (Tab. 1). The deforestation trend, characterized by a progressive decrease of the deforestation rate over the period 1990-2005 and a progressive increase of the deforestation after 2005, is valid for all four forest types except the spiny forest (Tab. 3). For the spiny forest, the deforestation rate during the period 2010-2013 was lower than on the period 2005-2010 (Tab. 3).

**Table 3:** Comparing Madagascar annual deforestation rates with previous studies on the period 1953-2013. Annual deforested areas (in thousands of hectares per year, Kha/yr) and annual deforestation rates (second number in parenthesis, in %/yr) are provided. For deforestation rates in %/yr, exact same numbers as in scientific articles and reports from previous studies (Harper *et al*., 2007; MEFT *et al*., 2009; ONE *et al*., 2015) have been reported. The way annual deforestation rates in %/yr have been computed in these previous studies can slightly differ from one study to another, but estimates always correct for the potential presences of clouds on satellite images and unclassified areas on forest maps. Annual deforested areas in Kha/yr have been recomputed from forest cover estimates in Tab. 2 (except for Harper *et al*. (2007) for the periods 1973-1990 and 1990-2000 for which annual deforested areas in Kha/yr were derived from numbers reported in the original publication, see methods) and do not correct for the potential presence of clouds.

| Forest type | Source | 1953-1973 | 1973-1990 | 1990-2000 | 2000-2005 | 2005-2010 | 2010-2013 |
| --- | --- | --- | --- | --- | --- | --- | --- |
| Total | Harper2007 | 91 (0.3) | 200 (1.7) | 81 (0.9) | - | - | - |
|  | MEFT2009 | - | - | 97 (0.8) | 53 (0.5) | - | - |
|  | ONE2015 | - | - | - | - | 95 (1.2) | 164 (1.5) |
|  | this study | 86 (0.6) | 205 (1.6) | 88 (0.9) | 42 (0.4) | 70 (0.7) | 90 (1.0) |
| Moist | Harper2007 | 94 (0.6) | 87 (1.7) | 32 (0.8) | - | - | - |
|  | MEFT2009 | - | - | 48 (0.8) | 17 (0.4) | - | - |
|  | ONE2015 | - | - | - | - | 20 (0.5) | 37 (0.9) |
|  | this study | 79 (1.0) | 101 (1.6) | 40 (0.8) | 21 (0.4) | 27 (0.6) | 54 (1.2) |
| Dry | Harper2007 | 11 (0.2) | 77 (1.9) | 20 (0.7) | - | - | - |
|  | MEFT2009 | - | - | 24 (0.7) | 11 (0.4) | - | - |
|  | ONE2015 | - | - | - | - | 51 (1.8) | 97 (2.3) |
|  | this study | 16 (0.4) | 71 (1.9) | 28 (0.9) | 12 (0.4) | 29 (1.0) | 31 (1.1) |
| Spiny | Harper2007 | -3 (-0.1) | 36 (1.2) | 28 (1.2) | - | - | - |
|  | MEFT2009 | - | - | 25 (1.2) | 23 (1.2) | - | - |
|  | ONE2015 | - | - | - | - | 25 (1.7) | 31 (1.7) |
|  | this study | -6 (-0.2) | 31 (1.3) | 20 (1.0) | 9 (0.5) | 13 (0.7) | 4 (0.3) |
| Mangroves | Harper2007 | - | - | 1 (0.2) | - | - | - |
|  | MEFT2009 | - | - | - | - | - | - |
|  | ONE2015 | - | - | - | - | 0 (0.3) | 0 (0.2) |
|  | this study | -3 (-1.7) | 1 (0.6) | 0 (0.2) | 0 (0.0) | 0 (0.0) | 0 (0.0) |

### 3.2 Comparison with previous forest cover change studies in Madagascar

Forest cover maps provided by previous studies over Madagascar were not exhaustive (unclassified areas) due to the presence of clouds on satellite images used to produce such maps. In Harper *et al*. (2007), the maps of years 1990 and 2000 include 0.5 and 1.12 Mha of unknown cover type respectively. Proportions of unclassified areas are not reported in the two other existing studies at the national level by MEFT *et al*. (2009) and ONE *et al*. (2015). With our approach, we produced wall-to-wall forest cover change maps from 1990 to 2014 for the full territory of Madagascar (Fig. 2). This allowed us to produce more robust estimates of forest cover and deforestation rates over this period (Tab. 1). Our forest cover estimates over the period 1953-2013 (considering forest cover estimates at national level and by ecoregions for all the available dates) were well correlated (Pearson’s correlation coefficient = 0.99) to estimates from the three previous studies (Tab. 2) with a RMSE of 300,000 ha (6% of the mean forest cover of 4.8 Mha when considering all dates and forest types together). These small differences can be partly attributed to differences in ecoregion boundaries. Despite significant differences in deforestation estimates (Tab. 3), a similar deforestation trend was observed across studies with a decrease of deforestation rates over the period 1990-2005, followed by a progressive increase of the deforestation after 2005.

### 3.3 Variation of forest fragmentation over time

Forest fragmentation has progressively increased since 1953 in Madagascar. We observed a continuous decrease of the mean distance to forest edge from 1953 to 2014 in Madagascar. The mean distance to forest edge has decreased to about 300 m in 2014 while it was of about 1.5 km in 1973 (Fig. 3). Moreover, a large proportion (73%) of the forest was located at a distance greater than 100 m in 1973, while almost half of the forest (46%) is at a distance lower than 100 m from forest edge in 2014 (Fig. 3). The percentage of lowly fragmented forest in Madagascar has continuously decreased since 1953. The percentage of forest belonging to the lowly fragmented class has fallen from 57% in 1973 to 44% in 2014. In 2014, 22% of the forest belonged to the two highest fragmented forest classes (less than 40% of forest cover in the neighborhood) while only 15% of the forest belonged to these two fragmentation classes in 1973 (Tab. 4).

**Figure 3:**
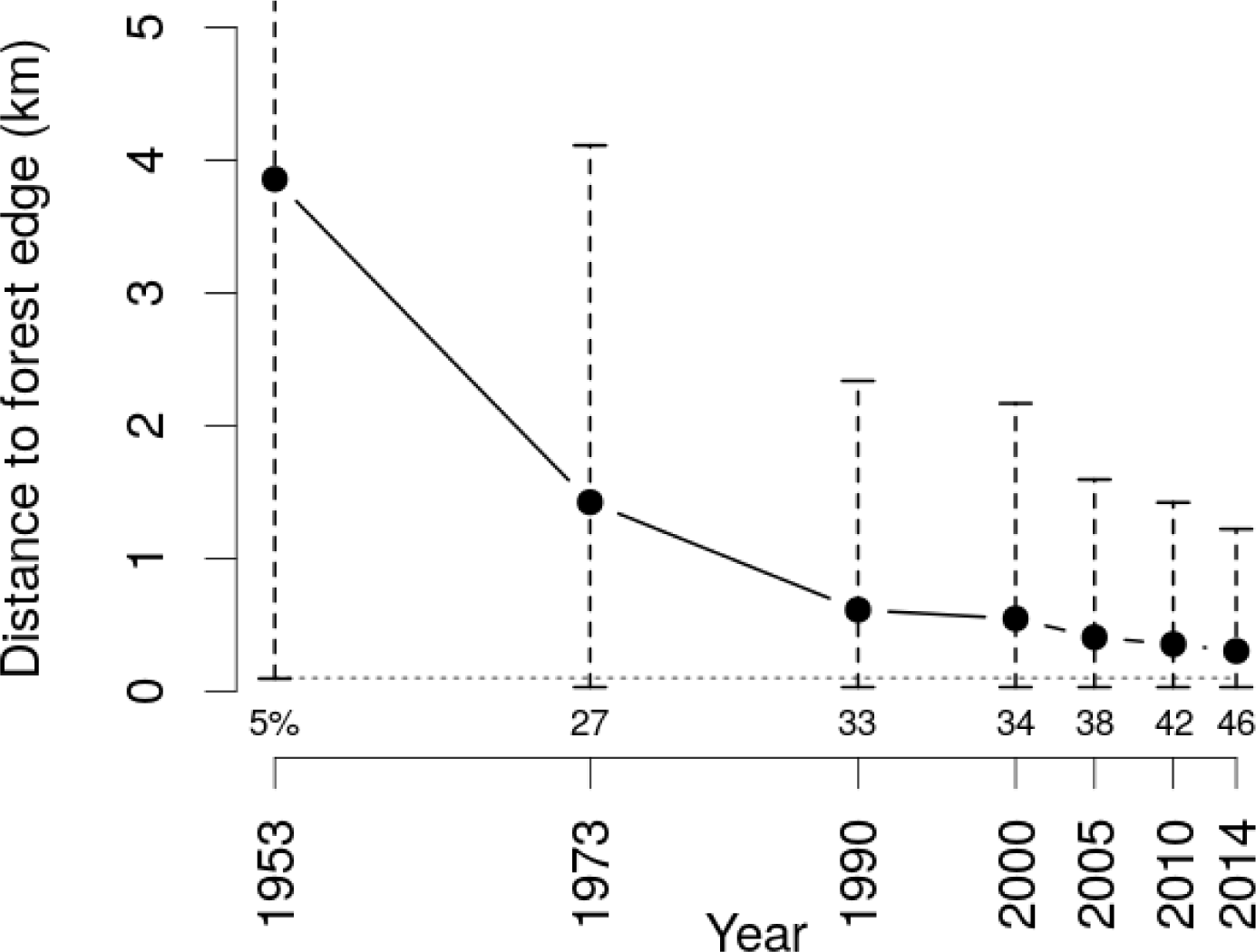
hange in distance to forest edge from 1953 to 2014 in Madagascar. Black dots represent the mean distance to forest edge for each year. Vertical dashed segments represent the 90% quantiles (5% and 95%) of the distance to forest edge. Horizontal dashed grey line indicates a distance to forest edge of 100 m. Numbers at the bottom of each vertical segments are the percentage of forest at a distance to forest edge lower than 100 m for each year.

**Table 4:**
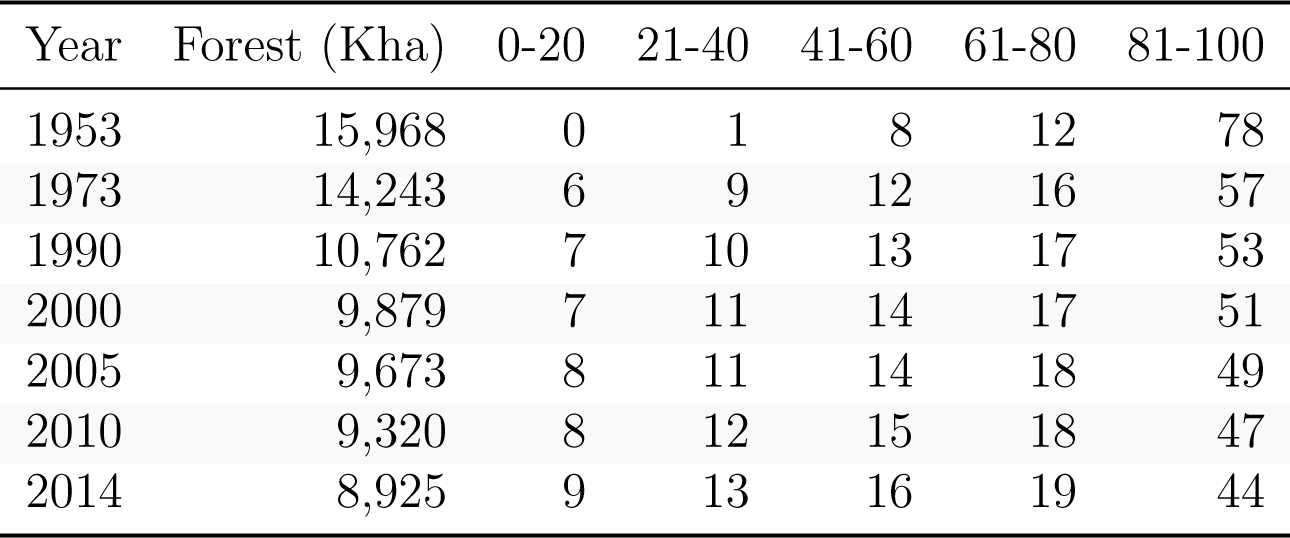
Change in forest fragmentation from 1953 to 2014 in Madagascar. Five forest fragmentation classes, based on the percentage of forest in the neighborhood, are defined: 0-20% (highly fragmented), 21-40%, 41-60%, 61-80% and 81-100% (lowly fragmented). The percentage of forest falling in each forest fragmentation class is reported for each year. Forest areas are provided in thousands of hectares (Kha).

| Year | Forest (Kha) | 0-20 | 21-40 | 41-60 | 61-80 | 81-100 |
| --- | --- | --- | --- | --- | --- | --- |
| 1953 | 15,968 | 0 | 1 | 8 | 12 | 78 |
| 1973 | 14,243 | 6 | 9 | 12 | 16 | 57 |
| 1990 | 10,762 | 7 | 10 | 13 | 17 | 53 |
| 2000 | 9,879 | 7 | 11 | 14 | 17 | 51 |
| 2005 | 9,673 | 8 | 11 | 14 | 18 | 49 |
| 2010 | 9,320 | 8 | 12 | 15 | 18 | 47 |
| 2014 | 8,925 | 9 | 13 | 16 | 19 | 44 |

## 4 Discussion

### 4.1 Advantages of combining recent global annual tree cover loss data with historical national forest cover maps

In this study, we combined recent (2001-2014) global annual tree cover loss data (Hansen *et al*., 2013) with historical (1953-2000) national forest cover maps (Harper *et al*., 2007) to look at six decades (1953-2014) of deforestation and forest fragmentation in Madagascar. We produced annual forest cover maps at 30 m resolution covering Madagascar for the period 2000 to 2014. Our study extends the forest cover monitoring on a six decades period (from 1953 to 2014) while harmonizing the data from previous studies (Harper *et al*., 2007; MEFT *et al*., 2009; ONE *et al*., 2015). We propose a generic approach to solve the problem of forest definition which is needed to transform the 2000 global tree cover dataset from Hansen *et al*. (2013) into a forest/non-forest map (Tropek *et al*., 2014). We propose the use of an historical national forest cover map, based on a national forest definition, as a forest cover mask. This approach could be easily extended to other tropical regions or countries for which an accurate forest cover map is available at any date within the period 2000-2014 (but preferably at the beginning of the period to profit from the full record of tree cover loss and derive long-term estimates of deforestation). For example, forest cover maps are available at 20 m resolution for Cameroon and Central African Republic for years 2000 and 2010 (Gross *et al*., 2017). When high resolution forest cover maps are not available, coarser resolution forest cover maps (leading to coarser deforestation estimates) could be extracted from global land cover products such as GLC2000 at 1 km resolution (Bartholomé & Belward, 2005) or CCI Land Cover at 300 m resolution (Li *et al*., 2018). Moreover, this approach could be repeated in the future with the release of updated tree cover loss data. We have made the **R**/GRASS code used for this study freely available in a GitHub repository (see Data availability statement) to facilitate application to other study areas or repeat the analysis in the future for Madagascar.

The accuracy of the derived forest cover change maps depends directly on the accuracy of the historical forest cover maps and the tree cover loss dataset. Using visual-interpretation of aerial images in 342 areas distributed among all forest types, *Harper et al*. (2007) estimated an overall 89.5% accuracy in identifying forest/non-forest classes for the year 2000. The accuracy assessment of the tree cover loss dataset for the tropical biome reported 13% of false positives and 16.9% of false negatives (see Tab. S5 in *Hansen et al*. 2013). These numbers rise at 20.7% and 20.6% respectively for the subtropical biome. In the subtropical biome, the lower density tree cover canopy makes it difficult to detect change from tree cover to bare ground. For six countries in Central Africa, with a majority of moist dense forest, Verhegghen *et al*. (2016) have compared deforestation estimates derived from the global tree cover loss dataset (Hansen *et al*., 2013) with results derived from semi-automated supervised classification of Landsat satellite images (Achard *et al*., 2014) and they found a good agreement between the two sets of estimates. Therefore, our forest cover change maps after 2000 might be more accurate for the dense moist forest than for the dry and spiny forest. In another study assessing the accuracy of the tree cover loss product across the tropics (Tyukavina *et al*., 2015), authors reported 4% of false positives and 48% of false negatives in Sub-Saharan Africa. They showed that 85% of missing loss occurred on the edges of other loss patches. This means that tree cover loss might be underestimated in Sub-Saharan Africa, probably due to the prevalence of small-scale disturbance which is hard to map at 30 m, but that areas of large-scale deforestation are well identified and spatial variability of the deforestation is well represented. A proper accuracy assessment of our forest cover change maps should be performed to better estimate the uncertainty surrounding our forest cover change estimates in Madagascar from year 2000 (Olofsson *et al*., 2014, 2013). Despite this limitation, we have shown that the deforestation trend we observed for Madagascar, with a doubling deforestation on the period 2010-2014 compared to 2000-2005, was consistent with the other studies at the national scale (MEFT *et al*., 2009; ONE *et al*., 2015).

Consistent with Harper *et al*. (2007), we did not consider potential forest regrowth in Madagascar (although Hansen *et al*. (2013) provided a tree cover gains layer for the period 2001-2013) for several reasons. First, the tree gain layer of Hansen *et al*. (2013) includes and catches more easily tree plantations than natural forest regrowth (Tropek *et al*., 2014). Second, there is little evidence of natural forest regeneration in Madagascar (Grouzis *et al*., 2001; Harper *et al*., 2007). This can be explained by several ecological processes following burning practice such as soil erosion (Grinand *et al*., 2017) and reduced seed bank due to fire and soil loss (Grouzis *et al*., 2001). Moreover, in areas where forest regeneration is ecologically possible, young forest regrowth are more easily re-burnt for agriculture and pasture. Third, young secondary forests provide more limited ecosystem services compared to old-growth natural forests in terms of biodiversity and carbon storage (Martin *et al*., 2013).

### 4.2 Natural forest cover change in Madagascar from 1953 to 2014

We estimated that natural forest in Madagascar covers 8.9 Mha in 2014 (corresponding to 15% of the country) and that Madagascar has lost 44% of its natural forest since 1953 (37% since 1973). If there are no doubts about the direct causes of deforestation in Madagascar, attributable to human activities such as slash-and-burn agriculture and pasture (Scales, 2011), there is ongoing scientific debate about the extent of the “original” forest cover in Madagascar, and the extent to which humans have altered the natural forest landscapes since their large-scale settlement around 800 CE (Burns *et al*., 2016; Cox *et al*., 2012). Early French naturalists stated that the full island was originally covered by forest (Humbert, 1927; Perrier de La Bâthie, 1921), leading to the common statement that 90% of the natural forests have disappeared since the arrival of humans on the island (Kull, 2000). More recent studies counter-balanced that point of view saying that extensive areas of grassland existed in Madagascar long before human arrival and were determined by climate, natural grazing and other natural factors (Virah-Sawmy, 2009; Vorontsova *et al*., 2016). Other authors have questioned the entire narrative of extensive alteration of the landscape by early human activity which, through legislation, has severe consequences on local people (Klein, 2002; Kull, 2000). Whatever the original proportion of natural forests and grasslands in Madagascar, our results demonstrate that human activities since the 1950s have profoundly impacted the natural tropical forests and that conservation and development programs in Madagascar have failed to stop deforestation in the recent years. Deforestation has strong consequences on biodiversity and carbon emissions in Madagascar. Around 90% of Madagascar’s species are forest dependent (Allnutt *et al*., 2008; Goodman & Benstead, 2005). Based on occurrence data for 2243 plant and invertebrate species, *Allnutt et al*. (2008) estimated that deforestation between 1953 and 2000 has led to an extinction of 9% of the species. The additional deforestation we observed over the period 2000-2014 (around 1 Mha of natural forest) worsen this result. Regarding carbon emissions, using the 2010 aboveground forest carbon map by Vieilledent *et al*. (2016), we estimated that deforestation on the period 2010-2014 has led to 40.2 Mt C of carbon emissions in the atmosphere (10 Mt C /yr) and that the remaining aboveground forest carbon stock in 2014 is 832.8 Mt C. Associated to deforestation, we showed that the remaining forests of Madagascar are highly fragmented with 46% of the forest being at less than 100 m of the forest edge. Small forest fragments do not allow to maintain viable populations and “edge effects” at forest/non-forest interfaces have impacts on both carbon emissions (Brinck *et al*., 2017) and biodiversity loss (Gibson *et al*., 2013; Murcia, 1995).

### 4.3 Deforestation trend and impacts on conservation and development policies

In our study, we have shown that the progressive decrease of the deforestation rate on the period 1990-2005 was followed by a continuous increase in the deforestation rate on the period 2005-2014. In particular, we showed that deforestation rate has more than doubled on the period 2010-2014 compared to 2000-2005. Our results are supported by previous studies (Harper *et al*., 2007; MEFT *et al*., 2009; ONE *et al*., 2015) despite differences in the methodologies regarding (i) forest definition (associated to independent visual interpretations of observation polygons to train the classifier), (ii) classification algorithms, (iii) deforestation rate computation method, and (iv) correction for the presence of clouds. Our deforestation rate estimates from 1990 to 2014 have been computed from wall-to-wall maps at 30 m resolution and can be considered more accurate in comparison with estimates from these previous studies. Our natural forest cover and deforestation rate estimates can be used as source of information for the next FAO Forest Resources Assessment (Keenan *et al*., 2015). Current rates of deforestation can also be used to build reference scenarios for deforestation in Madagascar and contribute to the implementation of deforestation mitigation activities in the framework of REDD+ (Olander *et al*., 2008).

The increase of deforestation rates after 2005 can be explained by population growth and political instability in the country. Nearly 90% of Madagascar’s population relies on biomass for their daily energy needs (Minten *et al*., 2013) and the link between population size and deforestation has previously been demonstrated in Madagascar (Gorenflo *et al*., 2011; Vieilledent *et al*., 2013). With a mean demographic growth rate of about 2.8%/yr and a population which has increased from 16 to 24 million people on the period 2000-2015 (United Nations, 2015), the increasing demand in wood-fuel and space for agriculture is likely to explain the increase in deforestation rates. The political crisis of 2009 (Ploch & Cook, 2012), followed by several years of political instability and weak governance could also explain the increase in the deforestation rate observed on the period 2005-2014 (see Smith *et al*., 2003 for a discussion on the link between governance and forest cover loss). These results show that despite the conservation policy in Madagascar (Freudenberger, 2010), deforestation has dramatically increased at the national level since 2005. Results of this study, including recent spatially explicit forest cover change maps and forest cover estimates, should help implement new conservation strategies to save Madagascar natural tropical forests and their unique biodiversity.

## 5 Author’s contribution

All authors conceived the ideas and designed the methodology; GV analysed the data and wrote the **R**/GRASS script; GV drafted the manuscript. All authors contributed critically to the drafts and gave final approval for publication.

## 6 Acknowledgements

The authors thank two anonymous reviewers for useful comments on a previous version of the manuscript. They also thank Peter Vogt for useful advice on which metric to use to estimate forest fragmentation. This study is part of the CIRAD’s BioSceneMada project (https://bioscenemada.cirad.fr) and the Joint Research Center’s ReCaREDD project (http://forobs.jrc.ec.europa.eu/recaredd). The BioSceneMada project is funded by FRB (Fondation pour la Recherche sur la Biodiversité) and the FFEM (Fond Fran¸cais pour l’Environnement Mondial) under the project agreement AAP-SCEN-2013 I. The ReCaREDD project is funded by the European Commission. The authors declare that there are no conflicts of interest related to this article.

## 7 Data accessibility

Data and code used for this study have been made permanently and publicly available on the CIRAD Dataverse repository so that the results are entirely reproducible:

- Input data: http://dx.doi.org/10.18167/DVN1/2FP7LR
- Code: http://dx.doi.org/10.18167/DVN1/275TDF
- Output data: http://dx.doi.org/10.18167/DVN1/AUBRRC

## Supplementary material

**Figure A1:**
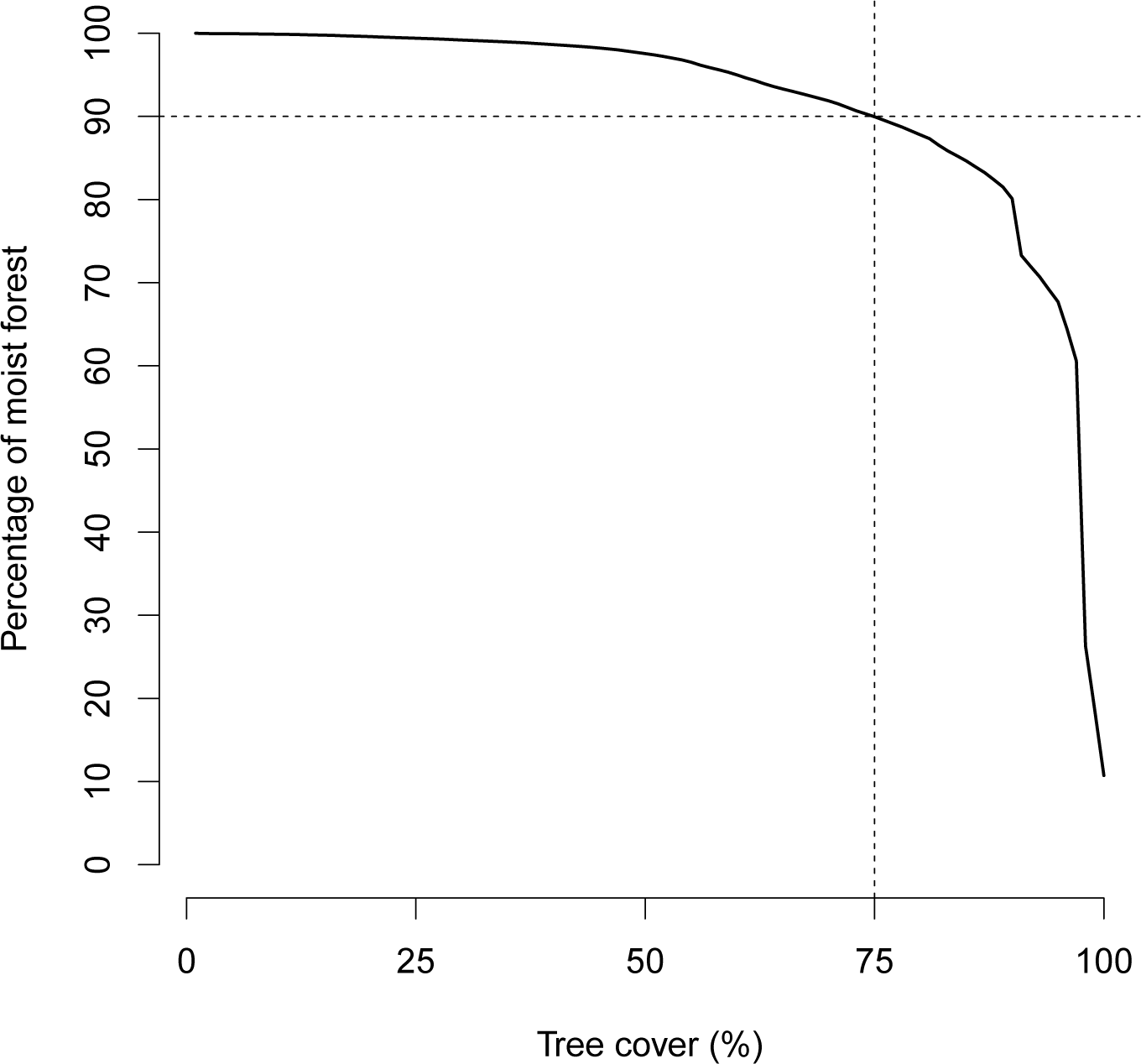
Selection of the tree cover threshold to define the land type for unclassified areas in 2000. We considered the forest in 2000 (Harper *et al*., 2007) in the moist ecoregion (see Fig. 1). We plotted the percentage of forest having a tree cover greater or equal to a given value specified by the x-axis. From this figure, we see that 90% of the moist forest in 2000 has a tree cover ≥ 75.

